# TabularQual: A spreadsheet-based format for annotating and curating logical models in SBML-qual

**DOI:** 10.64898/2026.05.31.727710

**Authors:** Luna Xingyu Li, Carissa Bleker, Sylvain Soliman, Laurence Calzone, Noriko Hiroi, Tomáš Helikar, Matthias König, Luiz Ladeira, Pedro T. Monteiro, Vincent Noël, Samuel Pastva, Albin Salazar, David Šafránek, Denis Thieffry, Eirini Tsirvouli, Anna Niarakis, John Gennari

## Abstract

Logical models are widely used to study regulatory and signaling systems, yet their reuse, annotation, and exchange across tools remain challenging. Although SBML Level 3 Qualitative Models (SBML-qual) provides a standard representation, its XML-based syntax is difficult to inspect and edit directly. Here we introduce TabularQual, a spreadsheet-based, community-driven representation for Boolean and multi-valued logical models, together with a bidirectional converter between spreadsheets and SBML-qual. The converter is accessible programmatically and via a web interface to support diverse user workflows. We further describe two integration strategies that enable existing modeling tools to operate with TabularQual, either through SBML-qual exchange or via direct support. Case studies using the Stress Knowledge Map and CaSQ demonstrate how this integration supports model construction, curation, and reuse. TabularQual provides a practical bridge between human-readable model representations and standardized executable formats, supporting reproducibility, interoperability, and community-driven model development.

## Introduction

Computational models are essential for formalizing biological mechanisms, simulating diverse system dynamics, and generating testable hypotheses. Among these, logical models are widely used to describe qualitative dynamics of gene regulatory and signaling networks, especially when quantitative kinetic data are scarce. As the number of published models and supporting tools increases, challenges in annotation, reuse, and interoperability become more pronounced (1,2).

To address these challenges, several formats have been developed for representing logical models. Among them, the Systems Biology Markup Language Qualitative Models (SBML-qual) has emerged as a central, community-maintained standard that facilitates model exchange and tool interoperability (3–5). SBML-qual supports structured annotations through RDF and provides an extensible representation of logical models; however, its XML-based syntax limits accessibility for routine inspection and editing. Many models remain poorly annotated, and annotation practices vary widely across groups. In practice, model metadata, node-level biological mappings, interaction descriptions, and transition logic are often captured in heterogeneous tabular formats, either as primary results, supplementary materials, or internal documentation.

For example, Fauré et al. use a table to describe a mammalian cell cycle model, in which ‘product’, ‘logical rules’, and ‘justification/references’ are presented as three columns for 10 species (6). In a more recent and larger model comprising 115 species, Selvaggio et al. describe the epithelial-to-mesenchymal transition process using a supplementary table with three tabs: ‘Node_annotations’, ‘Interactions_annotations’, and ‘Logical_rules’, each containing multiple columns to define attributes and provide annotations and comments (7). Furthermore, the Cell Collective, a large logical model repository and widely used modelling platform, provides a ‘knowledge base’ file for each model, which is essentially a table with multiple fields for annotations on ‘upstreamRegulator’, ‘regulatorSummary’, ‘references’, etc. (8).

The logical modeling community—through initiatives such as CoLoMoTo (9), COMBINE (10), SysMod (11), and BioModels (12)—has long advocated for improved annotation practices. Notably, recent community consensus (e.g., [BC]2 2021 workshop (2), [BC]2 2019 workshop (13), and CoLoMoTo/SysMod meetings) has emphasized the need for human-readable, tool-compatible formats to facilitate annotation, curation, and sharing.

In April 2025, during the HARMONY workshop (14), we presented a community-driven proposal: the use of standardized spreadsheets as an intuitive, extensible, and round-trip-compatible interface for annotating SBML-qual models. We argue that the tabular representation could improve accessibility for logical models, enabling direct editing, inspection, and validation without requiring familiarity with XML-based formats. This proposal received strong support. Attendees emphasized the dual role of such spreadsheets as: (i) a modeler-facing tool for model development and metadata documentation, and (ii) a curator-facing resource for repository submission and standardized annotation.

The approach builds on examples from well-known logical models (e.g., (6), (7), (15)), which all provide model annotations in a tabular format and align with efforts to automate conversion between spreadsheet formats and SBML-qual files. Community feedback from the following sessions further highlighted the importance of balancing minimalism (for adoption) and standardization (for interoperability), recommending features such as: support for multiple annotations per entity (via row-based structures); flexible identifier use (e.g., UniProt, Gene Ontology, PubMed); use of qualifiers to describe relationships between identifiers and model entities, including provenance tracking (e.g., isVersionOf, isDescribedBy); and support for automated tools (e.g., validation and identifier lookup).

In parallel, the broader systems biology community continues to push toward FAIR principles for Findable, Accessible, Interoperable, and Reusable models (16), and CURE principles for Credible, Understandable, Reproducible, and Extensible models (17). The spreadsheet-based annotations offer a pragmatic step toward achieving these goals for logical modeling.

This work formalizes the spreadsheet schema and annotation levels required to support this vision. It is intended as a starting point for community review, tool development, and eventual integration into major model repositories.

## Results

### The spreadsheet format

#### Design principles and scope

The spreadsheet format is designed as a lightweight, human-editable interface for annotating and curating logical models encoded in SBML-qual. It does not replace SBML-qual as an executable format, but instead provides an intermediate representation that exposes model structure, logic, annotations, and provenance in a form that is accessible to both modelers and curators.

Several design principles guided the development of the format. First, the representation prioritizes human readability and accessibility, allowing users to inspect, edit, and annotate models using familiar tabular interfaces. Second, the format enforces explicit semantics for model components, logical rules, and annotations, enabling deterministic conversion to and from SBML-qual. Finally, the format is designed to be extensible, allowing additional annotation layers or columns to be introduced in the future.

The scope of this initial version focuses on logical regulatory models represented in SBML-qual, including Boolean and multi-valued species, regulatory interactions, and transition logic. Representation of Petri nets and probabilistic Boolean networks is out of scope for this version.

#### Organization of the spreadsheet

The spreadsheet is organized into a set of structured sheets or tables, where each row represents a distinct model entity and each column corresponds to a defined attribute or annotation (except for the ‘Model’ sheet, see below). At minimum, three core sheets are required to define a valid logical model, ‘Model’, ‘Species’, and ‘Transitions’, with an optional ‘Interactions’ sheet for capturing pairwise relationships. Each core sheet includes the required fields needed to construct an SBML-qual model, along with optional fields for biological annotation, provenance, and documentation. The standard is designed to be extensible: additional sheets may be introduced in a fit-for-purpose manner to support specific use cases, and commonly adopted extensions could be formalized into the specification as community needs evolve. A complete specification and examples are provided in the Supplementary Material.

**Figure 1** illustrates the organization of the TabularQual format using part of an example model, which covers Boolean and multi-valued models (for two examples, see Supplementary Files S2, S3). In this representation, species correspond to the model components or nodes, transitions encode the logical update rules governing target species, and interactions capture optional pairwise regulatory relationships. Together, the four sheets provide complementary views of the same model, separating model metadata, species annotations, regulatory logic, and interaction evidence into a structured but linked tabular representation.

**Figure 1:**
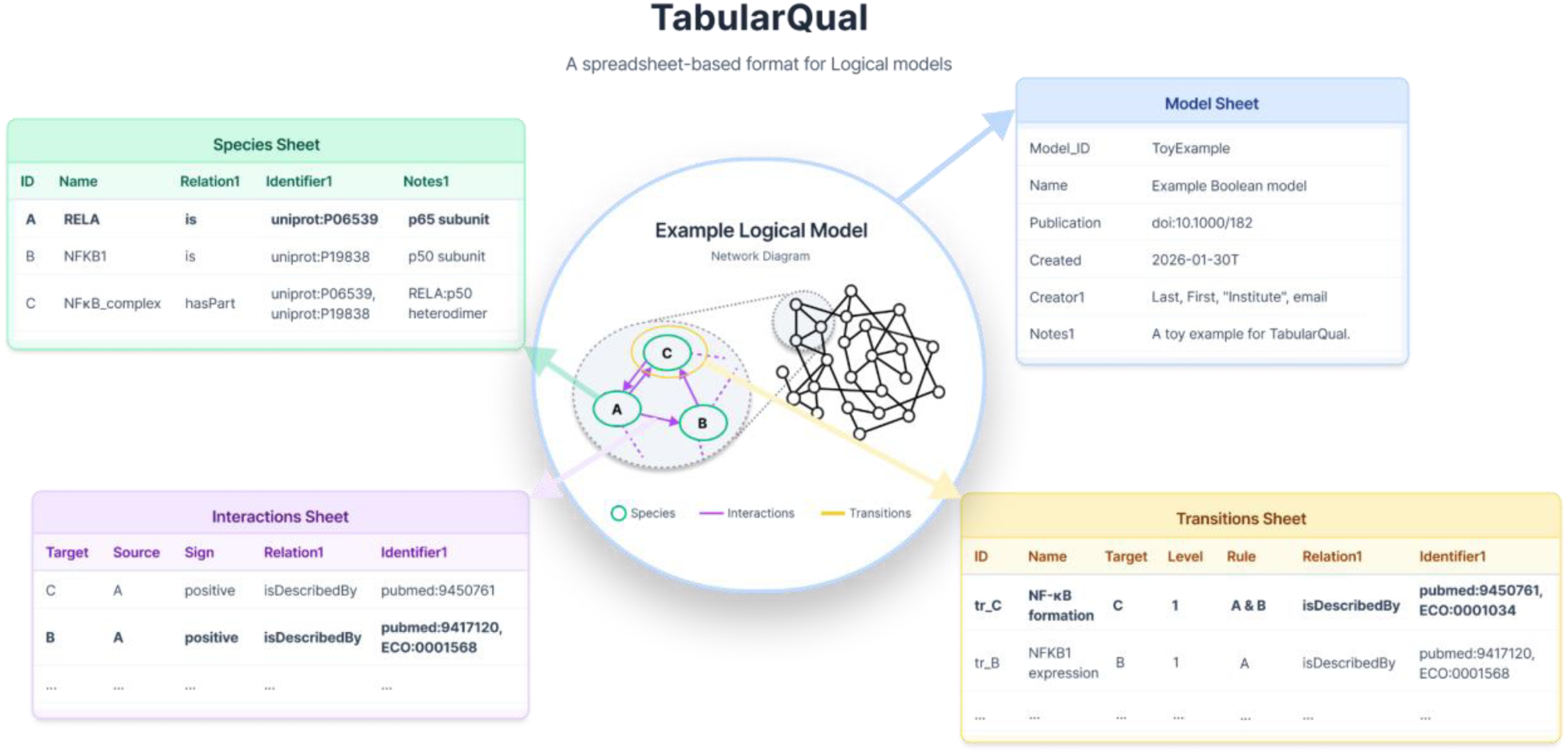
Example TabularQual spreadsheet representation of a logical model. Each worksheet corresponds to a class of model entities (Model, Species, Transitions, and Interactions).

The **Model** sheet captures metadata describing the model as a whole and provides provenance and contextual information, such as model source, biological process, organism, references, and contributor information. These fields map either to SBML model attributes or to RDF annotations using standard model qualifiers. Free-text notes are exported to SBML, whereas informal comments are retained only at the spreadsheet level.

The **Species** sheet defines all qualitative species (nodes). Each row corresponds to a single `QualitativeSpecies` element in SBML-qual and must include either an ID or a name. Optional fields specify attributes such as compartment, type (input, internal, or output), whether the species is constant, its initial level, and its maximum level (default 1 for Boolean models). Annotations are provided using compact identifiers (e.g., UniProt or NCBI Gene accessions) together with qualifiers that describe the relationship between the species and the referenced biological entity (e.g., ‘is’ or ‘isDescribedBy’). Multiple annotations per species are supported through comma-separated identifiers for the same qualifier or indexed columns for different qualifiers.

The **Transitions** sheet encodes the model’s regulatory logic. Each row defines a logical transition governing the state of a single target species and must include a `Target` species and a logical `Rule`. In multi-valued models, there may be multiple rows per target, i.e. one for each activating `Level`. Logical rules are expressed using a concise textual syntax with Boolean operators, e.g., ‘&’ (AND), ‘|’ (OR), ‘!’ NOT, ‘^’ (XOR). In addition, parentheses are used for grouping, and extended operators are used for multi-valued models, which can be deterministically mapped to SBML-qual function terms. Transitions may also be annotated with supporting evidence and provenance using the same identifier–qualifier format.

The optional **Interactions** sheet records evidence and provenance for pairwise regulatory influences between species independently of full logical rules. Each row specifies a `Source` species, a `Target` species, and the `Sign` of regulation. While interaction annotations are not required for model execution, they provide useful biological context, support graphical visualization, and can be leveraged by tools to assist in rule construction or validation.

Across all sheets, annotations follow a consistent semantic model based on RDF-style triples, using standard models and biology qualifiers. Identifiers are recommended to follow the compact ‘prefix:accession’ pattern registered with identifiers.org, but additional URLs are also supported to accommodate resources not covered by existing registries. This explicit representation of annotation semantics underpins reliable round-trip conversion between the spreadsheet format and SBML-qual.

The TabularQual standard defines a schema independent of file format. In practice, models can be represented either as multi-sheet spreadsheet files (e.g., XLSX) or as collections of delimited text files (CSV), depending on user workflows and tool requirements. Current implementations support both formats for bidirectional conversion.

### Implementation

TabularQual is designed to support multiple levels of integration with existing logical modeling tools, reflecting differences in tool architecture and user workflows. In the following section, we describe the two approaches for adopting the TabularQual format in existing tools, each with concrete examples.

#### The TabularQual converter between spreadsheet and SBML-qual

The most straightforward integration strategy is to use SBML-qual as a common exchange format. In this setting, the TabularQual converter serves as an intermediary layer that bridges spreadsheets and the SBML-qual workflow for bidirectional exchange.

The TabularQual converter enables round-trip transformation between the spreadsheet format and SBML Level 3 Qualitative Models. The converter is implemented in Python and relies on an in-memory representation of qualitative species, transitions, interactions, and logical rules for both conversion directions. As a first step, the tool will check and validate the TabularQual format for required fields and data types, while the SBML-qual format and annotation validation are supported via sbmlutils (18).

The converter is accessible via a command-line interface, a web interface, and a Python API, distributed as an installable package on PyPI for programmatic integration into external tools and workflows. The command-line interface supports scripted and reproducible workflows, while the web application provides an interactive environment for users without programming experience. The web interface is publicly available at https://tabularqual.streamlit.app/ and supports conversion in both directions, option selection, file preview, and direct download of results. Figure 2 illustrates the user interface for spreadsheet-to-SBML conversion.

**Figure 2:**
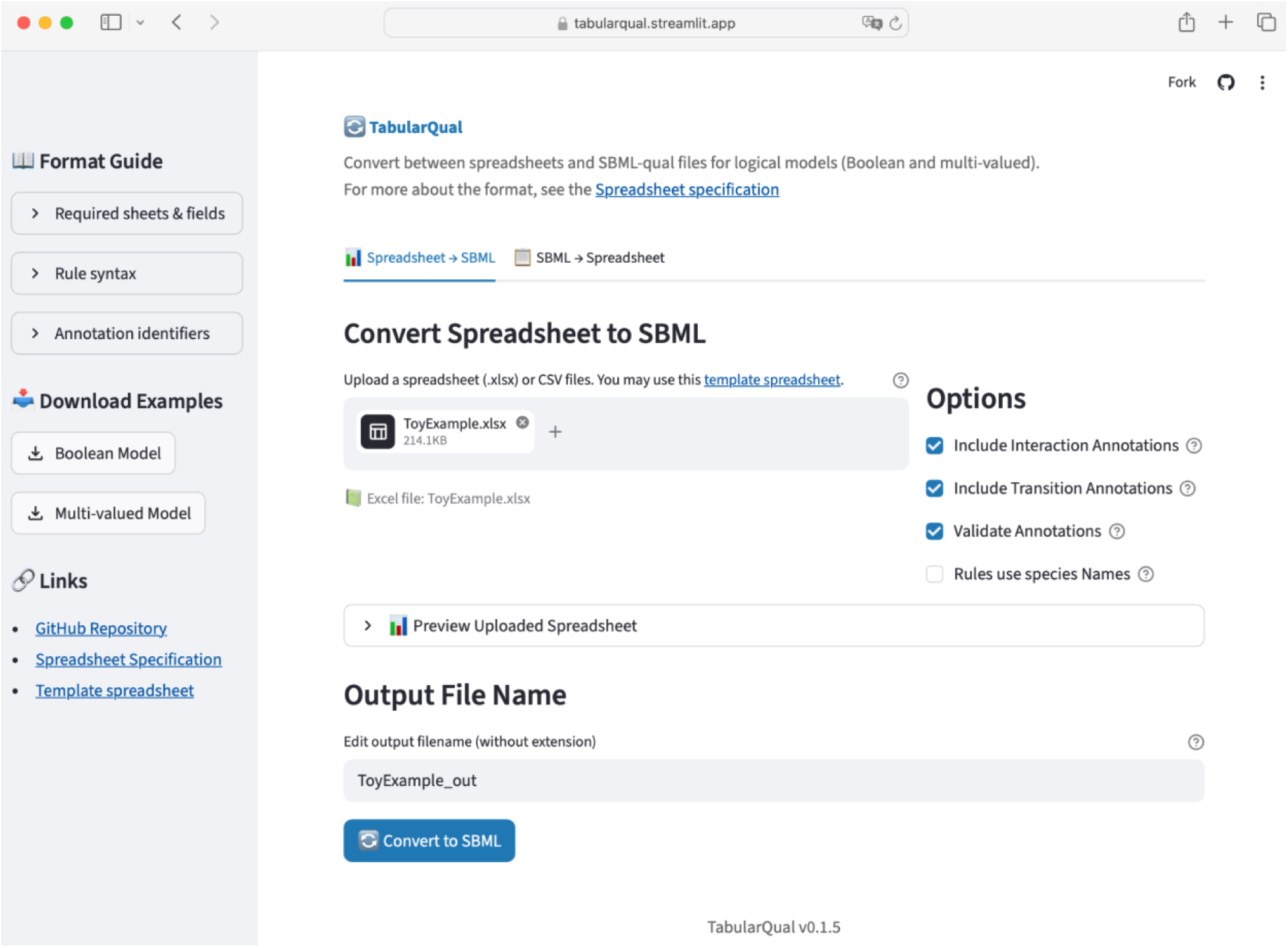
Web interface of the TabularQual round-trip conversion tool, showing the spreadsheet-to-SBML direction.

Both Boolean and multi-valued models are supported by the converter, with several configurable options to control rule representation, identifier usage, and output format. Logical rules may be expressed using species names instead of SBML identifiers to improve readability. To minimize information loss during round-trip conversion, the converter preserves model structure and logic, including qualitative species, transitions, input–output relationships, and, where supported, annotations. Users may further control whether annotations are retained at the interaction level, transition level, or both. By relying on SBML-qual as the exchange format, this design supports iterative spreadsheet-based curation while maintaining compatibility with existing logical modeling tools.

We evaluated the converter on 205 SBML-qual models in BioDivine Boolean Models Database (19), which were collected from multiple major repositories (e.g., GINsim, Cell Collective, BioModels) and individual publications. The original models without curation were used, and their model scale and sources can be found in Figure S1. This includes 154 Boolean models and 51 multi-valued models. Using a round-trip protocol (SBML → spreadsheet → SBML), 197 models achieved full fidelity (Figure 3, detailed in Figure S2). Topology and logical rules were exactly preserved across all 205 models, and annotation retention exceeded 95% for valid qualifier types. The remaining mismatches were due to legacy annotations containing commas. The average conversion time was below 0.1 seconds per model in each direction.

**Figure 3.**
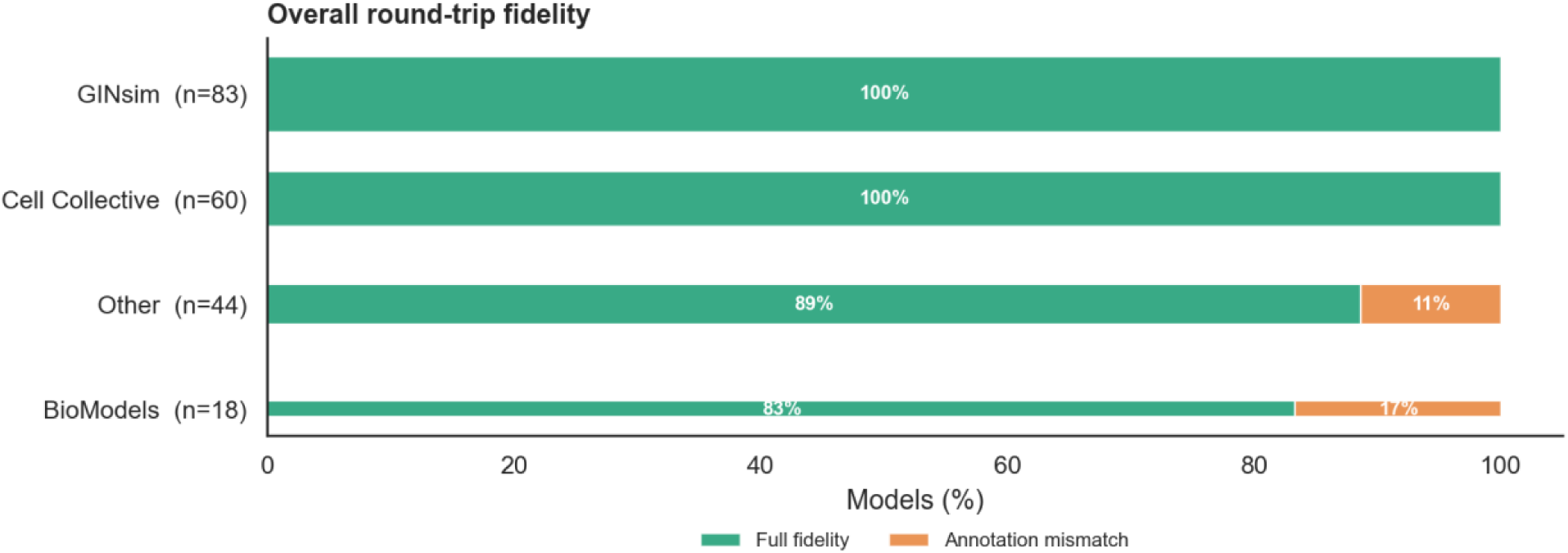
Round-trip conversion fidelity across 205 SBML-qual models. Overall round-trip fidelity by model source. Each bar shows the proportion of models achieving full fidelity (green) or annotation mismatch (orange). The height of the bars indicates the number of models from each source.

#### Compatibility with existing tools

Because the TabularQual converter supports bidirectional conversion with SBML-qual, any tool that imports or exports SBML-qual can indirectly operate on TabularQual representations. This typically requires a two-step process: users first employ the TabularQual converter to transform the spreadsheet into an SBML-qual file, which can then be loaded directly into these platforms for simulation, analysis, or visualization. Conversely, SBML-qual models exported from these tools can be converted back into spreadsheets for inspection and annotation.

**Table 1** summarizes the compatibility status of six commonly used logical modeling tools. We categorize them based on whether they require the use of the TabularQual converter (Section 3.1.1) or offer direct integration with the format (Section 3.2). Below, we provide brief summaries of each tool; note that we also provide additional use case details for the CaSQ and BoolDog systems.

**Table 1:** Logical modeling software tested to be compatible with TabularQual.

| Software | Technology | Supported formats | Functions | Compatibility | Link |
| --- | --- | --- | --- | --- | --- |
| AEON.py | Python | SBML-qual, AEON, BoolNet, etc. | Topological analysis and dynamic analysis | Uses TabularQual tool | https://github.com/sybila/biodivine-aeon-py |
| BoolDog | Python | SBML-qual, BoolNet, Primes | Dynamic analysis | Direct integration | https://github.com/NIB-SI/BoolDog/ |
| CasQ | Python | SBML-qual, CellDesigner, SBGN-ML | Boolean model construction and inference | Direct integration | https://gitlab.inria.fr/soliman/casq |
| CellCollective | JavaScript | SBML-qual | Topological analysis and dynamic analysis | Uses TabularQual tool | https://cellcollective.org/ |
| GINsim | Java | SBML-qual, GINML, BoolNet, Truth table, etc. | Multi-valued model definition, model annotation, topological analysis and dynamic analysis | Uses TabularQual tool | https://ginsim.github.io/ |
| MaBoSS | C++ | SBML-qual, MaBoSS, GINML, BoolNet | Dynamic analysis | Direct integration | https://maboss.curie.fr/ |

AEON.py is a Python library for symbolic analysis of asynchronous Boolean networks, supporting attractor analysis and formal verification (20). It accepts SBML-qual and related formats; therefore, TabularQual spreadsheet models can be converted to SBML-qual and loaded into AEON without loss of meaning. In addition, AEON.py supports partially specified networks and uncertainty in update rules, which aligns with the extensibility of the tabular format for representing incomplete or evolving biological knowledge.

BoolDog, a Python package for the synchronous analysis and semi-quantitative simulations of Boolean networks (21), can natively import TabularQual files into a Boolean model by internally converting TabularQual spreadsheets to supported SBML-qual. As we describe below, it is easier to review and modify model elements in a TabularQual spreadsheet than in the raw SBML-qual format.

CaSQ (CellDesigner as SBML-qual) enables the conversion of curated pathway diagrams into executable logical models (22). In addition to SBML-qual, CaSQ now supports direct export of TabularQual CSV files, allowing users to move seamlessly from graphical model construction to spreadsheet-based curation. As demonstrated in the CaSQ use case below, this facilitates transparent editing, annotation, and validation of logical rules.

Cell Collective is an interactive platform for constructing and simulating logical models of biological networks (8). It enables users to define regulatory relationships without requiring explicit mathematical formulation, while supporting dynamic simulation in a graphical, user-friendly environment. TabularQual is compatible with such workflows by providing a structured representation of models that aligns closely with the underlying abstractions used in Cell Collective. Through SBML-qual conversion, TabularQual models can be imported into Cell Collective for simulation and visualization, while existing models can be exported and converted back for annotation and curation.

GINsim (Gene Interaction Network simulation) is a Java software tool dedicated to the graphical construction, simulation, and topological and dynamic analysis of Boolean and multi-valued logical models (23). It supports synchronous and asynchronous update schemes, among others. GINsim relies on the jsbml library to import and export SBML-qual files. GINsim was used for round-trip tests, starting both from TabularQual spreadsheets and from GINsim, to ensure the preservation of the model’s qualitative behavior. In particular, the preservation of multi-valued logical rules and annotations were tested with this tool. GINsim also provides a comprehensive model repository of Boolean and multi-valued logical models (https://ginsim.github.io/models/). To promote the adoption of the TabularQual format, the GINsim team decided to provide copies of the TabularQual version for all models in their repository.

MaBoSS is a C++ software that simulates populations of asynchronous cells using continuous-time Markov processes over Boolean networks (24). MaBoSS can import any SBML-qual file, including those generated by the TabularQual spreadsheet. Moreover, pyMaBoSS (MaBoSS Python bindings) and WebMaBoSS (25), the online version of MaBoSS, can directly import the TabularQual spreadsheet. Furthermore, MaBoSS was one of the tools used to validate the preserved qualitative behavior of the models generated via the round-trip protocol.

This compatibility across tools demonstrates that TabularQual complements rather than replaces SBML-qual and existing model editing tools. The spreadsheet format serves as an accessible representation for model development, modification and documentation, while SBML-qual remains the canonical exchange format for tool interoperability. A practical adoption pathway is thereby provided for the broader community, allowing users to benefit from spreadsheet-based workflows while continuing to rely on established logical modeling software.

### Use cases of TabularQual

Beyond leveraging the converter, some tools benefit from direct coupling with the TabularQual spreadsheet format. This enables more seamless workflows for model construction, annotation, and curation, particularly for tools that already emphasize knowledge sources or graphical model design. In the following sections, we describe two examples of direct integration—Stress Knowledge Map and CaSQ—each illustrated by a concrete use case that demonstrates direct spreadsheet export and downstream interoperability.

#### Integration with Stress Knowledge Map

The Stress Knowledge Map (SKM) is a resource for molecular interactions in plants (26). A component of SKM, the plant stress signaling model (PSS), is a knowledge graph representation of plant stress responses, comprising stress perception, signaling and regulation, and protective processes. A contribution interface allows users to continuously add new reactions to PSS based on literature curation. Automated exports from the PSS (or parts thereof) to diverse formats (including interaction networks, SBML, and BoolNet) allow utilization of the PSS across analysis approaches. However, due to incomplete biological knowledge, heterogeneous curation practices, and the automated rule generation, exported models often need considerable curation before being ready to use in modeling investigations.

A driving design motivation for TabularQual is to ease the curation and extensions of models. Thus, we implemented a TabularQual export of PSS (or parts thereof) alongside the existing export formats available in the SKM web application. This format allows users to easily correct and curate dynamic models of plant stress signaling, in a format directly supported by the previously mentioned logical modeling tools.

As a use case, we interactively prepared a model of drought stress in the PSS Explorer (Figure 4A) and downloaded the model in TabularQual format. The model contains three connected pathways that trigger stomatal closure after drought. As downloaded, only one pathway was activated, due to common issues relating to the automated model extraction (entities in multiple compartments, inconsistent protein activation states). The spreadsheet format eased the identification and correction of these issues (Figure 4B), without needing to decipher and curate unannotated plain text (in BoolNet format). For example, one can immediately identify species names across multiple compartments or protein activation states within the *Species Sheet.* The corrected model was imported into BoolDog (21) and visualized via Cytoscape (27) (Figure 4C). The impact of drought on stomatal closure was simulated in BoolDog, which shows that all three pathways are correctly activated, resulting in the activation of the “Stomatal closure” process.

**Figure 4:**
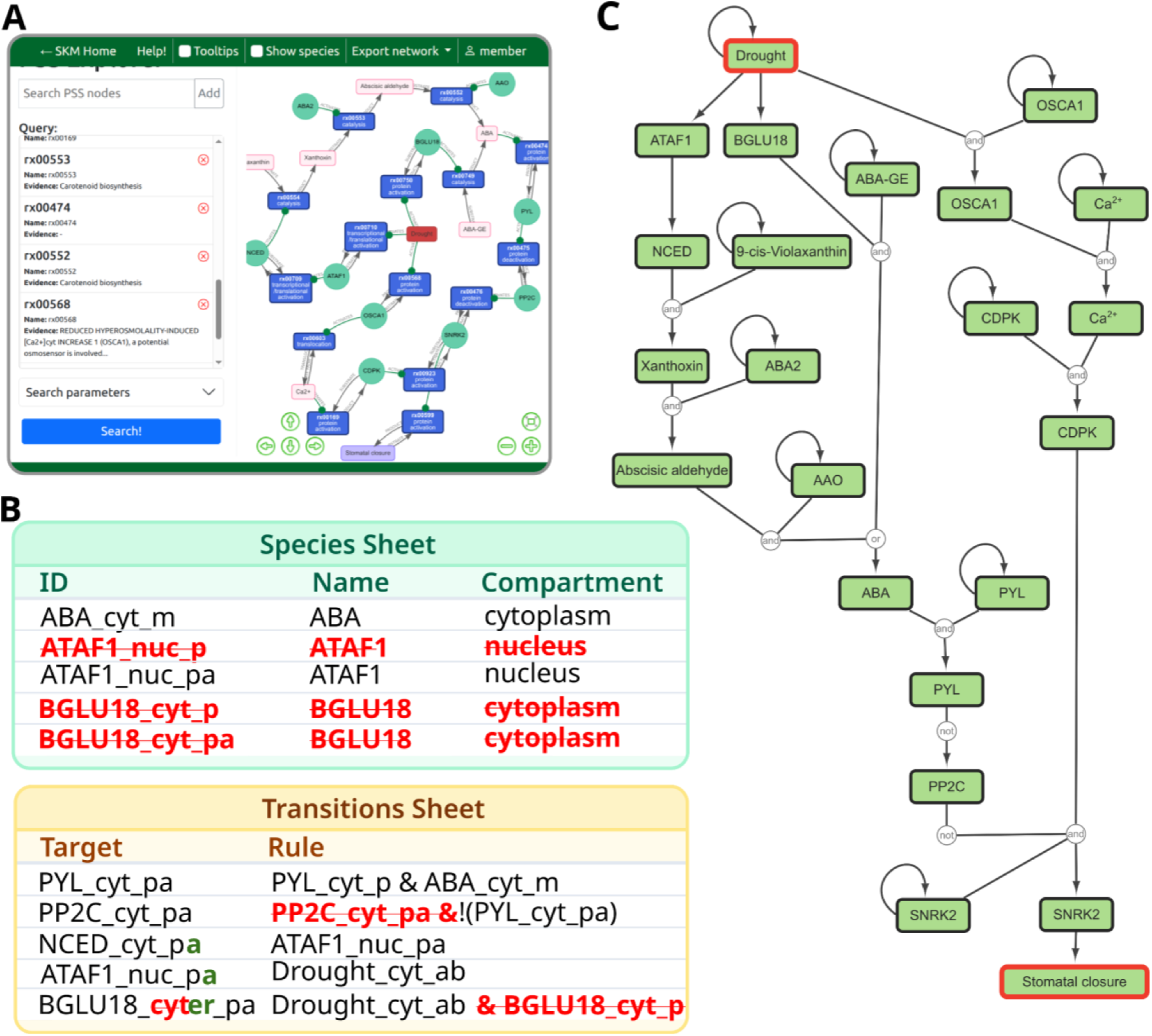
Stress Knowledge Map case study. (A) Building the model in the online PSS Explorer. (B) Corrections to the model in TabularQual format (updates in bold, red – removal, green – additions). (C) The corrected model visualized in Cytoscape using BoolDog.

#### Integration with CaSQ

CaSQ is a tool that infers dynamic Boolean models from static molecular interaction maps encoded in CellDesigner XML or, more recently, SBGN-ML (Process Description) formats (22). As molecular interaction maps can include hundreds to thousands of species, CaSQ was one of the first tools to help generate large-scale Boolean models in SBML-qual format, prompting the adoption of the format by many Boolean model analysis software, such as CellCollective (8) or WebMaBoSS (25).

Since support for the SBML-qual format was limited in existing tools at the beginning, CaSQ has always proposed generating multiple CSV files with information in a format more easily understood by its users. It was therefore natural to adapt this CSV generation to directly produce Spreadsheet SBML-qual CSV files, one file per sheet of the format proposed in this work. Currently, CaSQ generates the Model, Species and Transitions sheets. We decided not to export the (optional) interactions sheet, since CaSQ already exports these as a `.sif` file for Cytoscape.

In the example below, we show how to generate the spreadsheet files for an SBML-qual model using a CellDesigner XML as input. We have selected the apoptosis pathway from the COVID-19 Disease Map project (28). Diagrams can be downloaded in different formats; CaSQ can only handle CellDesigner XML or SBGN-ML. Next, using CaSQ, the static network can be transformed into an executable Boolean model (Figure 5A). Using the argument `-c`, five files are generated per entry. The SBML-qual file, the `.bnet` file, and three CSV files that include Models, Species, and Transitions information. The three CSV files correspond to the TabularQual spreadsheet, as obtained when the option to use species names in the Transitions sheet is selected.

**Figure 5:**
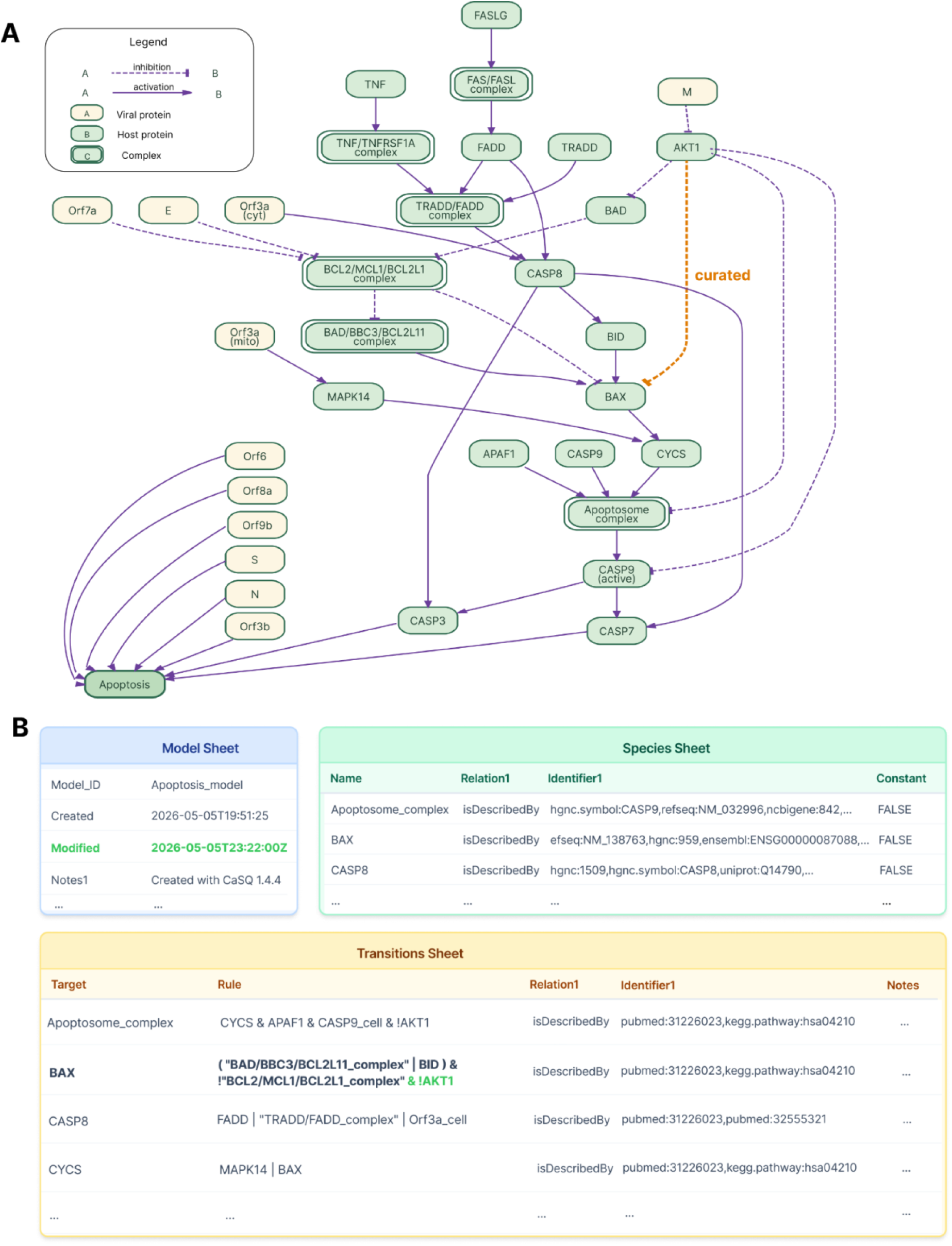
CaSQ case study. (A) Diagram of the apoptosis model generated by CaSQ. (B) Representation of the model in the TabularQual format, showing updates in bold, green represents additions.

One of the most important features of the TabularQual tool is that it allows the user to create a new model (by changing species or transitions) without opening any SBML-qual editor. Modifying the CSV files and creating an SBML-qual file from the Spreadsheets provides an alternative, quick way to modify and edit existing Boolean models.

In our example file, the apoptosis pathway model depicts both the intrinsic and extrinsic pathways: In the current model, BAX activation (which permeabilizes the mitochondrial membrane and triggers the intrinsic apoptosis cascade) is regulated by the following **Activators: BID, BAD/BBC3/BCL2L11_complex;** and **Inhibitor: BCL2/BCLXL/MCL1 complex**. AKT (activated downstream of PI3K survival signals) phosphorylates BAX at Ser184, retaining BAX in the cytosol in its inactive conformation and preventing its translocation to the mitochondrial outer membrane (29,30). This is distinct from the BCL-2 family proteins, which sequester BH3-only activators or directly bind BAX at the outer membrane. In this model, AKT1 already inhibits the Apoptosome (“!AKT1” in the Apoptosome and CASP9_cell_active rules), but not BAX directly. We propose adding a direct inhibition of BAX by AKT1 to better represent these biological mechanisms.

The rule can be modified directly in the Transition file, and the three CSV files (Species, Model, and modified_Transition) can then be used with TabularQual to obtain the modified SBML-qual model (Figure 5B). Model files, including both original and modified versions, are provided as Supplementary file S5.

Together, these two use cases demonstrate that TabularQual functions as an effective intermediate representation bridging model generation, curation, and execution. The SKM example highlights how spreadsheet-based editing facilitates the identification and correction of inconsistencies arising from automated model extraction. The CaSQ example illustrates complementary functionality, where large-scale models derived from graphical maps can be directly exported into TabularQual and refined without interacting directly with SBML syntax. Across both scenarios, TabularQual improves transparency of logical rules, supports structured annotation, and maintains compatibility with existing tools.

## Discussion

TabularQual addresses a practical gap between human-oriented model curation and machine-oriented model exchange. While SBML-qual remains the canonical executable representation for logical models, its XML syntax is not convenient for routine inspection, editing, or annotation. TabularQual provides a structured, human-readable layer over SBML-qual that preserves compatibility with existing standards while making model content easier to curate and review.

A central strength of this work is that the standard emerges from broader community discussion around recurring needs in logical modelling. We established this standard in response to long-recognized challenges in annotation consistency, interoperability, and curation workflows. The format builds on prior community efforts that called for minimum standards for annotation and improved reuse of logical models, and translates those goals into a concrete, operational schema for use in practice (2, 13). In this sense, TabularQual is both a technical contribution and a community standardization effort, intended to support communication among modelers, curators, repositories, and software developers.

An important contribution of TabularQual is the explicit organization of several annotation layers within one coherent schema, including model-level metadata, species-level annotations, transition logic, and optional interaction-level evidence. By separating these layers while preserving links among them, the format makes it easier to distinguish biological identity, model structure, evidence, and provenance. This is particularly useful for logical models, where the same biological system may be represented by multiple alternative rules, levels of abstraction, or curation histories.

To facilitate reuse and reliable round-trip conversion, users of TabularQual should follow a small number of **consistent practices**. Identifiers should be stable, unique, and as meaningful as possible for the readability of logical rules, while the human-readable name field can preserve preferred biological labels. Annotations should use appropriate qualifiers and identifiers. Species annotations should preferably refer to stable biological entities or processes using compact identifiers, whereas literature or provenance information should be attached using qualifiers such as ‘isDescribedBy’ or ‘isDerivedFrom’, depending on the intended meaning. In general, interaction-level annotations are most appropriate for evidence supporting individual pairwise regulations, whereas transition-level annotations are more suitable for evidence supporting a complete logical rule or a combined regulatory mechanism. Model-level metadata should also be recorded as completely as possible, including a model identifier, descriptive name, source, publication, organism, and relevant biological process, together with creator information, timestamps, version labels, and explanatory notes whenever available. Notes and comments should also be used distinctly: notes are appropriate for free-text content that should remain associated with the exported SBML model, whereas comments are best reserved for spreadsheet-only remarks and should not be relied upon to carry essential model content because they are ignored during conversion.

A second major contribution is the practical adoption pathway enabled by the converter and the integration strategies described in this work. Because TabularQual supports bidirectional conversion with SBML-qual, it can be smoothly incorporated into existing modelling workflows without requiring every tool to implement a new representation.

At the same time, direct support from tools such as CaSQ, MaBoSS, and BoolDog demonstrates that the format is not only theoretically compatible but also usable in real workflows. This dual strategy lowers the barrier to adoption. For users, it enables spreadsheet-based inspection and editing without sacrificing compatibility with existing standards. For tool developers, it offers both a lightweight exchange route through SBML-qual and a more direct route for native support when appropriate.

This work has several **limitations**. First, the current standard is intentionally scoped to logical regulatory models represented in SBML-qual. It supports Boolean and multi-valued models, but it does not aim to cover all qualitative or discrete formalisms. Petri nets, probabilistic Boolean networks, and hybrid extensions remain outside the scope of the present specification and would require additional conventions in future versions. Second, the core tables are not sufficient to enable fully reproducible modelling studies. In particular, simulation settings, analysis workflows, and execution environments are not part of TabularQual itself. For this reason, the format should be viewed as one component of a broader reproducibility stack (e.g., the CoLoMoTo rather than a complete solution on its own.

A practical consideration for dissemination is how to package the tabular representation. Because TabularQual is independent of any single serialization format, the same model can be represented either as one multi-sheet spreadsheet file, such as an XLSX workbook, or as a set of separate CSV files corresponding to the core tables. A combined spreadsheet file is often more convenient for manual inspection, editing, and sharing, since all sheets remain bundled in a single document and sheet-level structure is preserved. By contrast, separate CSV files are simpler for command-line workflows, version control, and integration with existing tools that already import or export delimited text files. The main limitation of the CSV representation is that the model is distributed across multiple files, making it easier to lose contextual coherence or separate one table from the others. In such cases, a COMBINE archive can provide a useful container for packaging the full set of CSV files together, optionally alongside related documentation or simulation settings, while preserving a single distributable research object (31).

More broadly, TabularQual aligns with ongoing efforts in systems biology to improve the accessibility, interoperability, and reusability of computational models. Its value lies not only in the design of the tables themselves, but in offering a shared interface between human-oriented curation and machine-oriented model exchange. By standardizing a spreadsheet-based representation that is readable, extensible, and compatible with SBML-qual, TabularQual provides a sustainable foundation for future tool development, repository workflows, and community-driven model curation. We expect that continued feedback from the logical modelling community will help refine the specification, extend support across tools, and further establish TabularQual as a practical standard for representing logical models. We will maintain and update the TabularQual converter and specification in response to community needs, to ensure long-term usability, compatibility, and continued adoption.

## Acknowledgement

We gratefully acknowledge the late Claudine Chaouiya for her leadership in developing SBML-qual and her foundational contributions to logical modeling. Her guidance and support were instrumental in shaping this work.

## Data availability

The TabularQual specification is provided as Supplementary File S1. The TabularQual converter is available as an open-source Python package through PyPI. Source code, documentation, example files, and scripts for reproducing the conversion workflows described in this manuscript are available at the project repository: https://github.com/sys-bio/TabularQual.

The SBML-qual models used for round-trip evaluation were obtained from the publicly available BioDivine Boolean Models Database at https://github.com/sybila/biodivine-boolean-models.

## List of abbreviations

API: application programming interface
CoLoMoTo: Consortium for Logical Models and Tools
COMBINE: COmputational Modeling in BIology Network
CSV: comma-separated values
CURE: Credible, Understandable, Reproducible, and Extensible
FAIR: Findable, Accessible, Interoperable, and Reusable
PSS: Plant Stress Signalling model
RDF: Resource Description Framework
SBML: Systems Biology Markup Language
SBML-qual: Systems Biology Markup Language Level 3 Qualitative Models package
SKM: Stress Knowledge Map
SysMod: Computational Modelling of Biological Systems
XLSX: Microsoft Excel Open XML spreadsheet format

