## Supplementary material for "TabularQual: A spreadsheet-based format for annotating and curating logical models in SBML-qual": TabularQual_Supplementary_File_S1_Specification.docx

TabularQual Specification

Version 1 Release 1

### 1. Introduction

**Logical models** are vital in systems biology for formalizing mechanisms and simulating dynamics. The systems biology markup language QualitativeModels (SBML-qual) was introduced to encode logical models as a standard format (1–3). However, the community still faces significant challenges regarding model annotation and interoperability (4,5). Many models remain poorly annotated, and annotation practices differ widely across groups. Historically, model metadata, node-level biological mappings, interaction descriptions, and transition logic have often been recorded in **ad hoc tabular formats**, which are provided as supplementary material of model publication, or just kept as an internal reference.

To address this, the logical modeling community—under initiatives such as CoLoMoTo (6), COMBINE (7), SysMod (8) and BioModels (9)—has long advocated for **better annotation practices**. Notably, recent community consensus (e.g., [BC]2 2021 workshop (4), [BC]2 2019 workshop (10) and CoLoMoTo/SysMod meetings) has emphasized the need for **human-readable, tool-compatible formats** to facilitate annotation, curation, and sharing.

In April 2025, during the HARMONY workshop series, we presented a community-driven proposal: the use of **standardized spreadsheets** as an intuitive, extensible, and round-trip-compatible interface for annotating SBML-qual models. This proposal received strong support. Attendees emphasized the dual role of such spreadsheets as:

- a **modeler-facing** tool for model development and metadata documentation,
- and a **curator-facing** resource for repository submission and standardized annotation.

The approach builds on examples from well-known logical models (e.g., Fauré 2006 (11), Selvaggio 2020 (12), Zerrouk 2024 (13)), which all provide model annotation in a tabular format and align with efforts to **automate conversion** between spreadsheet formats and SBML-qual files. Community feedback from the CALM session further highlighted the importance of balancing **minimalism** (for adoption) and **standardization** (for interoperability), recommending features such as:

- Support for multiple annotations per entity (via row-based structures);
- Flexible identifier use (e.g., UniProt, GO, PubMed);
- Use of qualifiers to describe relationships between identifiers and model entity, including provenance tracking (e.g., isVersionOf, isDescribedBy);
- And support for automated tools (e.g., validation and identifier lookup).

In parallel, the broader systems biology community continues to push toward FAIR principles—Findable, Accessible, Interoperable, and Reusable models (14), and CURE principles–Credible, Understandable, Reproducible, and Extensible (15). The spreadsheet-based annotations offer a pragmatic step toward achieving these goals for logical modeling.

This specification document formalizes the spreadsheet schema and annotation levels required to support this vision. It is intended as a starting point for community review, tool development, and eventual integration into major model repositories.

### 2. Technical specification

This section formally defines the spreadsheet-based annotation format for logical models. It describes the tabular structure, element attributes for species and transitions, recommended format for annotations, and intended mappings between spreadsheet entries and the SBML-qual elements.

Importantly, this specification is independent of the underlying file format. Implementations may use spreadsheet-based formats (e.g., XLSX), plain-text tabular formats (e.g., CSV), or other equivalent encodings.

The associated converter is available via the following repository: <https://github.com/sys-bio/TabularQual>.

#### 2.1 Data organization

The spreadsheet consists of several tables (sheets), where each **row** represents a distinct model entity (**species**, **transition**, or **interaction**). And each **column** corresponds to a defined attribute or annotation of that entity. In addition, a **model** sheet is used to provide other information about the model as a whole.

At a minimum, two core tables are required to define a logical model, with the last two being optional:

- **Species** sheet: Defines QualitativeSpecies (nodes), their attributes, and annotations.
- **Transitions** sheet: Defines transitions (logical rules), their attributes, and annotations.
- (Optional) **Model** sheet: Defines the model and its metadata.
- (Optional) **Interactions** sheet: Provides annotations of the interactions (each edge) between two species.

Each sheet contains one or two data fields (columns) that are required to define a model. In addition, optional columns for annotation can be used to store external references and provenance data. For example, species can be linked with their biological entity ID from UniProt or NCBI gene; transitions and interactions can be supported by evidence from literature, etc.

Additional sheets and columns may be included if needed, but they will not be considered as part of the standard nor be kept in the converted SBML-qual model. In the template, two additional sheets are provided:

- **README** sheet: Provides basic instructions for using the template.
- **Appendix** sheet: Describe the options for certain columns (see 2.2).

#### 2.2 Definitions of the tables

This section provides definitions of the fields in each table. The order of the tables can be changed, as long as the tab names and headers remain the same.

Typically, there is only one entry per cell; if multiple entries are provided, they should be comma-separated. For identifiers used for annotations, we recommend using compact identifiers that follow the [prefix]:[accession] pattern registered at [*identifiers.org*](http://identifiers.org) or [*bioregistry.io*](https://bioregistry.io/) when possible. For example, ‘uniprot:P19838’ may be used to refer to protein NFKB1 with identifier ‘P19838’ in UniProt, and ‘DOI:10.1093/bib/bbac212’ points to the publication with that DOI. Additional URLs are supported for other resources, e.g., ‘<https://www.uniprot.org/uniprotkb/P19838/>’.

##### 2.2.1 Model sheet

The model sheet contains general information about the model as a whole, including annotations of model provenance and metadata.

Table 1 provides the definition and an example entry of each field in the model sheet. In the Mapping column, blue represents the corresponding SBML attributes of model elements; green represents the qualifier used for annotations of the model object.

Table 1: Definition of the fields in the **Model** sheet.

| **Field** | **Mapping** | **Definition** | **Example** |
| --- | --- | --- | --- |
| **Model_source** | bqmodel:is | URL or accession of the model or the spreadsheet itself. | https://www.ebi.ac.uk/biomodels/MODEL2307180001 |
| **Model_ID** | model.id | Identifier for the model; must be a valid SBML *SId* (ASCII, no spaces). | ToyExample_boolean |
| **Name** | model.name | Descriptive label of the model, can be any text. | Example Boolean model of NF-κB signaling pathway |
| **Publication** | bqmodel:isDescribedBy | Primary publication(s) supporting the encoded model; compact IDs stored with a qualifier. | DOI:10.1093/bib/bbac212 |
| **Origin_publication** | bqmodel:isDerivedFrom | Paper(s) describing the biological system from which the present model was derived. | pubmed:38272919 |
| **Origin_model** | bqmodel:isDerivedFrom | Accession(s) of parent model in compact identifier. | biomodels.db:MODEL2307180001 |
| **Taxon** | bqbiol:hasTaxon | NCBI taxonomy compact ID of the species (animal) described by the model. | taxonomy:9606 |
| **Biological_process** | bqbiol:isVersionOf | Biological process(es) described in the model. | GO:0007249 |
| **Created** | dcterms:created | ISO-8601 timestamp when this model is created. | 2025-04-04T17:08:28Z |
| **Modified** | dcterms:modified | Last edit timestamp when the model is modified. | 2025-07-29T16:56:00Z |
| **Creator[1, 2, …]** | dcterms:creator | Information about the creator of the model: [family name, given name, "organization", email]; if more than one creator, add new rows with header “**Creator2**”, “**Creator3**”, … | Li, Luna, "Center for Reproducible Biomedical Modeling", |
| **Contributor[1, 2, …]** | dcterms:contributor | An additional person or organization that has contributed to the model; encoded like **Creator**. |  |
| **Version** | model.notes | Other versions of the model can be listed here using the file name. | ToyExample_multivalue.xlsx |
| **Notes[1, 2, …]** | model.notes | Free-text as *notes* in the SBML model. | “This is an example note.” |
| **Comments** |  | Informal comments NOT exported to SBML. |  |

##### 2.2.2 Species sheet

The Species sheet lists every *QualitativeSpecies* (node) that appears in the logical model; usually this refers to genes or gene products in biological processes.

Either **Species_ID** or **Name** is required for a valid SBML-qual model. When only one is provided, the other may be generated automatically; all other columns provide optional attributes or annotations.

Table 2 provides the definition and an example entry of each field in the species sheet. SBML-qual attributes (blue in the Mapping column) are written exactly as they appear in the specification with capitalization; annotation qualifiers (green) should follow the BioModels Qualifiers vocabulary.

Table 2: Definition of the fields in the **Species** sheet.

| **Field** | **Mapping** | **Definition** | **Example** |
| --- | --- | --- | --- |
| **Species_ID** | QualitativeSpecies.id | Identifier for the species; must be a valid SBML *SId* (ASCII, no spaces). | NFKB1 |
| **Name** | QualitativeSpecies.name | Descriptive label of the species. | NFκB protein p50 subunit |
| **Relation[1, 2, …]** | bqbiol qualifiers | Qualifier for identifier[1, 2, …], respectively; defaults to 'is' | is |
| **Identifier[1, 2, …]** |  | Compact identifiers for Relation[1, 2, …], respectively. | uniprot:P19838 |
| **Compartment** | QualitativeSpecies.compartment.id | The compartment in which the species is located; must be a valid SBML *SId* (ASCII, no spaces). | cytosol |
| **Type** |  | Describe the role of the species in a network; Choose from: Input, Internal, Output. | Internal |
| **Constant** | QualitativeSpecies.constant | Whether the level of the species is fixed or can be varied; Choose from: True, False. | False |
| **InitialLevel** | QualitativeSpecies.initialLevel | Initial value of the species; non-negative integer; cannot exceed MaxLevel if both are set. | 0 |
| **MaxLevel** | QualitativeSpecies.maxLevel | Maximum value of the species; non-negative integer; default ‘1’ for Boolean models. | 1 |
| **Notes[1, 2, …]** | QualitativeSpecies.notes | Free-text as *notes* for the species. | “p50 subunit of NF-κB complex” |
| **Comments** |  | Informal comments NOT exported to SBML. |  |

Because either **Species_ID** or **Name** is required in the species sheet, they can be used interchangeably in both Interactions and Transitions sheets. Names that conform to the SBML *SId* syntax may be used directly. Names that do not conform to *SId* requirements (e.g., containing spaces or slashes) must be enclosed in double quotes when used. During SBML export, names are resolved to their corresponding identifiers as defined in the Species sheet.

##### 2.2.3 Interactions sheet

The Interactions sheet records evidence and provenance for pairwise influences between species.

It is an **optional** sheet and purely for annotating pair-wise relations between nodes, rather than the transitions as a whole. In addition, softwares may use it to decorate graphical views or to auto-generate default Transition inputs/outputs.

Table 3: Definition of the fields in the **Interactions** sheet.

| **Field** | **Mapping** | **Definition** | **Example** |
| --- | --- | --- | --- |
| **Target** | Transition.Output.id (QualitativeSpecies.id) | Identifier for the regulated species; the output that participates in a transition. Must be an existing Species_ID. | A |
| **Source** | Transition.Input.id (QualitativeSpecies.id) | Identifier for the regulator species; the input that is affected by a transition. Must be an existing Species_ID. | D |
| **Sign** | Transition.Input.sign | Whether the contribution of this input is positive, negative, both (dual) or unknown. | positive |
| **Relation[1, 2, …]** | bqbiol qualifiers | Qualifier for identifier[1, 2, …], respectively; defaults to 'isDescribedBy' | isDescribedBy |
| **Identifier[1, 2, …]** |  | Compact identifiers for Relation[1, 2, …], respectively. | pubmed:20300203, ECO:0007392 |
| **Notes[1, 2, …]** | Transition.Input.notes | Free-text as *notes* for the interaction. | “IKK complex activates RELA by phosphorylating IκBα. ” |
| **Comments** |  | Informal comments NOT exported to SBML. |  |

##### 2.2.4 Transitions sheet

The transitions sheet provides core regulatory information of the model. For Boolean models, each row in the Transitions sheet defines the logical rule (*Transition*) that governs one output species; multi-valued species can take multiple rows to define the rules for each activation *Level*.

Only **Target** and **Rule** are required. Transition_ID and Name are optional and will be assigned a default value in the SBML model.

Table 4: Definition of the fields in the **Transitions** sheet.

| **Field** | **Mapping** | **Definition** | **Example** |
| --- | --- | --- | --- |
| **Transitions_ID** | Transition.id | Identifier for the transition; Optional, if provided, must be a valid SBML *SId* (ASCII, no spaces). | tr_A |
| **Name** | Transition.name | Descriptive label of the transition. | RELA activation |
| **Target** | Transition.Output.id (QualitativeSpecies.id) | Identifier for the regulated species; the output that participates in a transition. Must be an existing Species_ID. | A |
| **Level** | Transition.functionTerm.resultLevel | Effect of the transition on the corresponding target species; non-negative integer; default ‘1’ for Boolean models. | 1 |
| **Rule** | Transition.functionTerm.math | Logical rules of the transition; expressions using Boolean operators or integers for constant values. See Table 5. | A \| (D & !C) |
| **Relation[1, 2, …]** | bqbiol qualifiers | Qualifier for identifier[1, 2, …], respectively; defaults to 'isDescribedBy' | isDescribedBy |
| **Identifier[1, 2, …]** |  | Compact identifiers for Relation[1, 2, …], respectively. | pubmed:20300203, ECO:0007392 |
| **Notes[1, 2, …]** | Transition.notes | Free-text as *notes* for the interaction. | “IKK activates RELA by phosphorylating IκBα. ” |
| **Comments** |  | Informal comments NOT exported to SBML. |  |

The representation of logical rules uses a conventional expression, as in many popular logical modeling tools (e.g., GINsim, BoolNet), where the regulatory effect can be described as Boolean operations of regulators. In addition, a constant integer level (e.g., 0, 1, or k for multi-valued models) or a keyword FALSE/TRUE can be used to indicate that the target species is forced to the specified level. Table 5 gives a list of allowed symbols to be used in Boolean expressions.

Table 5: Symbols that can be used in **Transitions** - **Rule** field for Boolean expressions

| **Symbol** | **Definition** | **Example** |
| --- | --- | --- |
| & | Boolean “AND” operator | A & B |
| \| | Boolean “OR” operator | A \| C |
| ! | Boolean “NOT” operator | !A |
| ^ | Boolean “XOR” operator | A ^ B |
| () | Parentheses to prioritize regulation | A \| (D & !C) |
| = | Equality operator | A = 1 |
| TRUE**,** FALSE | Boolean constants; equivalent to 0 and 1 | TRUE |
| 0**,** 1 | Constant level assignment; forces the target species to the given level | 0 |
| “**,** ” | Used to enclose Species Names with special characters | IKK & ! "NFκB complex" |
| `space` | Spaces will be ignored in parsing the expression |  |

In addition, several other operators can be used for multi-valued rules, for example, use of comparison symbols to set the threshold for a given species.

Table 6: Additional symbols that can be used for multi-valued expressions

| **Symbol** | **Definition** | **Example** |
| --- | --- | --- |
| : | Used only in multi-valued models to represent the threshold level of an input (equal to >=) | A & B:2 |
| = | Equality operator | A = 2 |
| >**,** <**,** >=**,** <= | Used only in multi-valued models for comparison operators | A & (B >= 1) |
| 0**,** 1**,** 2**, …** | Constant level assignment; forces the target species to the given level | 2 |

When parentheses are absent, operators bind in the following order (highest to lowest): comparison operators (>=, >, <=,<, =, !=), then NOT (!), AND (&), XOR (^), OR (|). Comparison expressions such as A >= 2 are treated as atomic terms before any logical operator is applied. All binary operators are left-associative, so a chain of the same operator groups left-to-right. For example, ‘A & B | C’ is equivalent to ‘(A & B) | C’, and ‘A | B | C’ is equivalent to ‘(A | B) | C’. Parentheses can always be used to make grouping explicit and are recommended whenever the intended precedence might be ambiguous.

Examples:

- A & B - Both A and B are active (level ≥ 1 for multi-valued)
- A ^ B - Exactly one of A or B is active (XOR)
- A >= 2 | B < 1 - A is at level 2+ OR B is inactive
- (A & B) | (!C & D != 1) - Complex grouped expression
- CycE:2 & !p53 - CycE is greater than or equal to level 2 AND p53 inactive
- FALSE — target is forced inactive (level 0)
- 2 — target is forced to level 2 (multi-valued)

##### 2.2.5 Other sheets

- **README** – instructions on using the template. Content is ignored by converters but for user guidance only.
- **Appendix** – controlled vocabularies and allowed option lists referenced by Type, Constant, Relation, Sign.

Both sheets are informational only and may be removed from the spreadsheet.

### 3. Concepts used

This specification adopts a number of concepts and data types defined in the SBML Level 3 Core and SBML Level 3 Qualitative Models specifications. The most important concepts are summarized below for clarity and to support consistent usage across tabular representations and tools.

##### Systems Biology Markup Language QualitativeModels (SBML-qual)

SBML-qual is the standard format used to encode logical models, serving as the canonical executable representation for logical models in systems biology. TabularQual is developed based on the QualitativeModels package for SBML Level 3 Version 1.

**Reference**:<http://identifiers.org/combine.specifications/sbml.level-3.version-1.qual.version-1.release-1>

##### QualitativeSpecies (species or nodes)

A QualitativeSpecies is an element (node) that appears in the logical model, typically representing genes or gene products in biological processes. They are defined in the Species sheet, which records their attributes (such as Compartment, InitialLevel, and MaxLevel) and annotations.

**Example**: Species *C* refers to ‘*NFκB complex*’.

##### Transitions (logical rules)

A Transition defines the core regulatory information and the logical rule that governs one output species. For Boolean models, each row in the Transitions sheet defines the rule for a specific output species.

**Example**: ‘*C = A & B*’ describes that species *C* can be activated/expressed when species *A* and *B* are both active/present.

##### Interactions

Interactions record evidence and provenance for pairwise influences between species. The Interactions sheet is optional and used solely to annotate pairwise relations between nodes, separate from the transitions (logical rules) as a whole.

**Example**: The relationship between species *C* (target) and species *B* (source) is positive.

##### SBML identifiers (*SId*) and names

An *SId* is a string-based identifier with strict syntactic constraints, which has been used in **Model_ID**, **Species_ID**, **Transitions_ID**, and **Compartment** in the spreadsheet format. It follows the same definition as in SBML Level 3 Version 2 specification. Briefly, it has the following properties:

- It must start with a letter (A–Z, a–z) or underscore (_)
- It may contain only letters, digits (0–9), and underscores
- It is case-sensitive, and equality is determined by exact string matching
- It does not allow spaces, slashes, or other special characters
- It uses plain ASCII characters (no Unicode)

In contrast, the optional name attribute is intended solely for human readability. Names may contain spaces and arbitrary text, are not required to be unique, and must not be used for structural references within a model.

For improved readability, species names are allowed in **Transitions** - **Target**, **Rule** and **Interactions** - **Target**, **Source,** where names contain invalid symbols (e.g., \, (), or spaces) can be enclosed by double quotes (e.g., “NFκB protein”), while duplicate names are automatically disambiguated using numeric suffixes (_1, _2, …). This ensures SBML validity while allowing users to retain meaningful names in the spreadsheet.

**Examples**:

- SId: A, csa001, NFKB1
- Name: RELA, Inflammatory response, NFκB protein p50 subunit
- Rule using ID: A = csa001 & ! NFKB1
- Rule using name: RELA = “Inflammatory response” & ! “NFκB protein p50 subunit”

**Reference**: <https://identifiers.org/combine.specifications:sbml.level-3.version-2.core.release-2>

##### Compact identifier

A Compact Identifier is a unique string consisting of a Prefix (assigned by curator), a colon (‘:’), and an Accession (e.g., local identifier string). The Prefix is composed of an optional Provider Code, and an assigned Namespace, separated by a slash (‘/’).

They will be put as identifiers.org URLs in the SBML model using the following form:

https://identifiers.org/[provider_code/]namespace:accession

**Examples**: uniprot:P19838, [pubmed:22140103](https://identifiers.org/pubmed:22140103), [ec-code:1.1.1.1](https://identifiers.org/ec-code:1.1.1.1), [taxonomy:9606](https://identifiers.org/taxonomy:9606)

**Reference**: <https://docs.identifiers.org/pages/identification_scheme.html>

##### Qualifiers (bqmodel/bqbiol)

Qualifiers are used to describe the relation between a model component and the resource used to annotate it.

One can view the annotation of a model component as a statement in the form of a 'triple'. The resource used in the annotation is the 'object', while the qualifier is the 'predicate'. In the cases of the model qualifiers, the 'subject' of the relation is the modelling concept represented by the model component referenced by the annotation. The modelling concept may be the model itself, a mathematical construct, or a hypothesis that is proposed, changing the way we previously understood the model, etc. In the cases of the biology qualifiers, the 'subject' of the relation is the biological or biochemical object represented by the enclosing model element.

There are two kinds of qualifiers used for different purposes: 1) model qualifiers, and 2) biology qualifiers. Below is the list of qualifiers that may be used in a model:

**Model qualifiers**: is, isDerivedFrom, isDescribedBy, isInstanceOf, hasInstance

**Biology qualifiers**: is, hasVersion, isVersionOf, isDescribedBy, hasPart, isPartOf, hasProperty, isPropertyOf, encodes, isEncodedBy, isHomologTo, occursIn, hasTaxon, hasSource, hasSink, hasMediator, hasMultiplier, hasPhysicalEntity

**Reference**: <https://identifiers.org/combine.specifications:qualifiers-1.1>

### 4. Examples

- [ToyExample.xlsx](https://docs.google.com/spreadsheets/d/1_xY0VboBhejg8tWnGNyAFpEA_ZU762B0/edit?usp=sharing&ouid=105819375684543832411&rtpof=true&sd=true): A toy model in Boolean formalism.
- [ToyExample_multivalue.xlsx](https://docs.google.com/spreadsheets/d/1gcRtkNhJDny3R5ZtimwIIu-s5okNmp35/edit?usp=sharing&ouid=105819375684543832411&rtpof=true&sd=true): A multi-valued version of the toy model.
- [Faure2006.xlsx](https://docs.google.com/spreadsheets/d/1NIEpBQx7rZXizkUt6fk-lZquP1uS-vnH/edit?usp=drive_link&ouid=105819375684543832411&rtpof=true&sd=true): The Fauré 2006 model, widely used as an example in tools such as GINsim, Cell Collective, MaBoSS, BoolNet, etc.
  - [Faure2006.sbml](https://drive.google.com/file/d/1v8BVHoMAWCokhmiMHqBRwi1kbzwCHJ_i/view?usp=drive_link): Equivalent SBML file of the Fauré 2006 model.
- [ThieffryThomas1995_multivalue.xlsx](https://docs.google.com/spreadsheets/d/10Ce0TB1UztRDaEnRT8DnjoH3n3Jn7mt7/edit?usp=drive_link&ouid=105819375684543832411&rtpof=true&sd=true): A multi-valued model on phage lambda.
- [ThieffryThomas1995_operators.xlsx](https://docs.google.com/spreadsheets/d/1b_qYRnNNxoXBiwa65hF17YCiP_xkTj5r/edit?usp=drive_link&ouid=105819375684543832411&rtpof=true&sd=true): The same multi-valued model using a different rule syntax.
  - [ThieffryThomas1995_multivalue.sbml](https://drive.google.com/file/d/1k8oXKi_eYzxK8SLz-jUu58I5Zba16Aly/view?usp=drive_link): Equivalent SBML file of the Thieffry & Thomas 1995 model.

### 5. Best practices

To facilitate reuse and reliable round-trip conversion, the users of TabularQual should follow a small number of consistent practices. First, **identifiers** should be stable, unique, and as meaningful as possible for the readability of the logical rules. In addition, the human-readable name field can preserve the preferred biological label. See Appendix A for details of how unique IDs versus human-readable names are handled.

Second, **annotations** should use appropriate qualifiers and identifiers. Species annotations should preferably refer to stable biological entities or processes using compact identifiers, while literature or provenance information should be attached with qualifiers such as ‘isDescribedBy’ or ‘isDerivedFrom’, depending on the intended meaning. In general, interaction-level annotations are most appropriate for evidence about individual pairwise regulations, whereas transition-level annotations are more suitable when the evidence supports a complete logical rule or a combined regulatory mechanism. Consistent use of qualifiers is important to preserve the distinction between biological identity, provenance, and supporting evidence (4,5,10).

Third, **model-level metadata** should be recorded as completely as possible. At minimum, this includes a model identifier, a descriptive name, a source or origin, a publication, an organism, and a relevant biological process. Creator and contributor information, timestamps, version labels, and explanatory notes should also be included whenever available. Such metadata improves traceability, facilitates repository submission, and makes the model easier to interpret and update in the future (4,10,14,15).

Fourth, **notes and comments** should be used for different purposes. Notes fields are appropriate for free-text information that should remain associated with the exported SBML model, such as biological clarification, curation rationale, or warnings about interpretation. By contrast, Comments are best reserved for local spreadsheet-only remarks and should not be relied upon to carry essential model content, because they are ignored during conversion.

Finally, for dissemination and reproducible exchange, **TabularQual files** should ideally be distributed together with the corresponding SBML-qual model, related documentation, and, when available, simulation settings. Although this is not required by the spreadsheet standard itself, bundling these materials in a COMBINE archive provides a practical way to preserve context and reduce the risk of separating the model from its metadata, annotations, and auxiliary files (7,16).
