## Supplementary material for "TabularQual: A spreadsheet-based format for annotating and curating logical models in SBML-qual": readme.txt

This folder contains all the files used in the TabularQual - SKM case study.

TabularQual spreadsheet exported from PSS:
- tabqual-pss-drought_export.xlsx

Modified TabularQual file after curation:
- tabqual-pss-drought_curated.xlsx

SBML model file converted from the modified TabularQual files:
- tabqual-pss-drought_curated_out.sbml
