## Supplementary material for "TabularQual: A spreadsheet-based format for annotating and curating logical models in SBML-qual": readme.txt

This folder contains all the files used in the TabularQual - CaSQ case study.

CellDesigner XML obtained from the COVID-19 Disease Map:
- apoptosis.xml

Files generated by CaSQ using the CellDesigner XML:
- apoptosis.sbml
- apoptosis.bnet
- apoptosis.xgmml
- apoptosis_Model.csv
- apoptosis_Species.csv
- apoptosis_Transitions.csv
Note that the three csv files are TabularQual files.

Modified TabularQual files after curation:
- mod_Model.csv
- mod_Species.csv
- mod_Transitions.csv

SBML model file converted from the modified TabularQual files:
- mod_out.sbml
