## Supplementary material for "TabularQual: A spreadsheet-based format for annotating and curating logical models in SBML-qual": TabularQual_Supplementary_Information.docx

### Supplementary Files

#### Supplementary File S1. TabularQual specification document

This file provides the formal TabularQual specification. It defines the purpose and scope of the format, the organization of the core tables, the required and optional fields for model-level metadata, qualitative species, transitions, and interactions, and the conventions for representing semantic annotations and provenance. The specification is intended to support deterministic conversion between TabularQual and SBML-qual while remaining independent of a particular serialization format.

#### Supplementary File S2. Toy Boolean example

This file provides the Boolean toy model shown in Figure 1 of the manuscript. The workbook illustrates the standard TabularQual organization using the Model, Species, Transitions, and Interactions sheets. It is intended as a compact example for users and tool developers to inspect the expected spreadsheet structure and basic annotation conventions.

#### Supplementary File S3. Toy multi-valued example

This file provides a multi-valued version of the toy model. It demonstrates how TabularQual represents qualitative species with maximum levels greater than one and how transition rows can encode level-specific activation rules. The example complements Supplementary File S2 by showing the extension from Boolean to multi-valued logical models.

#### Supplementary File S4. Stress Knowledge Map case-study model files

This file contains the model files used for the Stress Knowledge Map case study. The files document the workflow from model export through spreadsheet-based curation and downstream use in logical-modeling tools. This case study illustrates how TabularQual facilitates manual inspection and correction of model inconsistencies arising from automated extraction of plant stress-signaling networks.

#### Supplementary File S5. CaSQ case-study model files

This file contains the model files used for the CaSQ case study, including the original and modified versions of the apoptosis-pathway model. The files support the example in which CaSQ-generated CSV files corresponding to TabularQual tables are edited directly and then converted back to SBML-qual. This case study demonstrates spreadsheet-based modification of logical rules without requiring users to edit SBML-qual XML directly.

### Supplementary Figures

#### Figure S1. Characteristics of the evaluated model dataset.

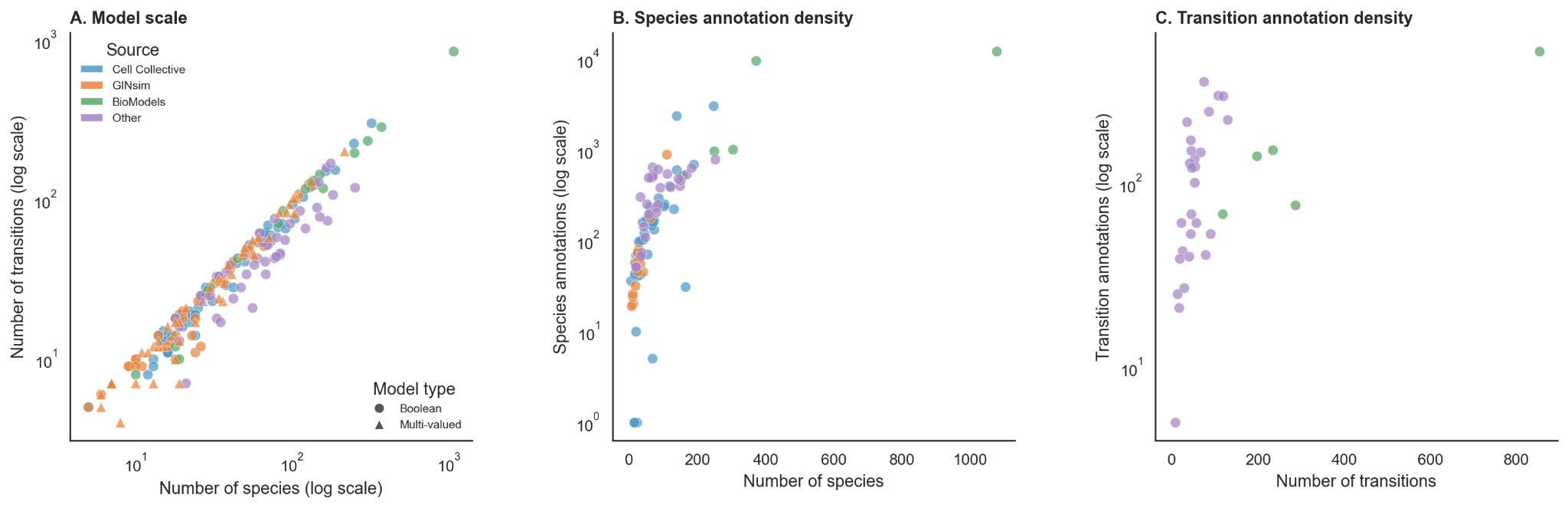

In all three graphs, color indicates the source of the model. (A) Model scale: number of transitions plotted against number of species, both on log scales, for each of the 205 evaluated models. The dataset includes 154 Boolean models, shown as circles, and 51 multi-valued models, shown as triangles. (B) Species annotation density: total number of species annotations, on a log scale, plotted against the number of species. Match or mismatch status indicates whether species counts were preserved after round-trip conversion. (C) Transition annotation density: total number of transition annotations, on a log scale, plotted against the number of transitions, with match or mismatch status as in (B).

#### Figure S2. Detailed round-trip conversion fidelity.

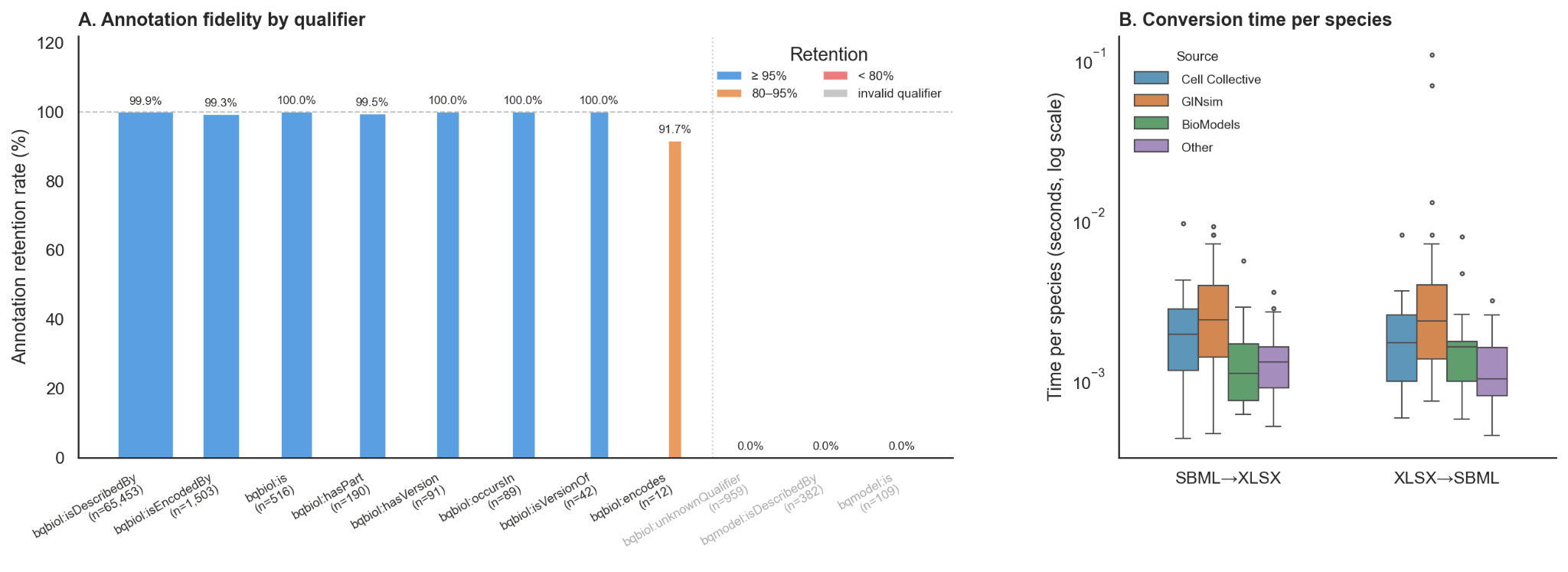

(A) Annotation retention rate by qualifier type. Bars show the percentage of annotation identifier pairs retained after round-trip conversion, colored by retention tier: blue, 95% or higher; orange, 80% to 95%; gray, invalid qualifier in the original model. (B) Conversion time per species, on a log scale, for both SBML-to-XLSX and XLSX-to-SBML conversion, grouped by model source.

In the BioDivine benchmark dataset, two models, 146_BUDDING-YEAST-FAURE-2009 and 159_BUDDING-YEAST-CORE, were conversion time outliers because their transition rules are written in fully expanded disjunctive normal form. These models contain rules with up to 650 and 513 OR clauses, respectively, compared with a dataset mean of 2.67 and median of 1 OR clause per rule. Their longest rules contain 76,318 and 56,075 characters, exceeding Excel’s 32,767-character cell limit and therefore requiring CSV output. The main bottleneck occurs during XLSX/CSV-to-SBML conversion, where each OR clause must be encoded as nested MathML <apply> elements. Across all 10,694 rules, only 10 exceed 100 OR clauses, with these two models accounting for most of them.
